# KSMoFinder - Knowledge graph embedding of proteins and motifs for predicting kinases of human phosphosites

**DOI:** 10.1101/2025.10.21.683733

**Authors:** Manju Anandakrishnan, Karen E. Ross, Chuming Chen, K. Vijay-Shanker, Cathy H. Wu

**Affiliations:** Center for Bioinformatics and Computational Biology, University of Delaware, Newark, United States of America; Department of Biochemistry and Molecular and Cellular Biology. Georgetown University Medical Center, Washington, DC, United States of America

## Abstract

**Motivation:** Protein kinases regulate cellular signaling pathways through a cascade of phosphorylation activity, selectively targeting specific residues on substrate proteins (phosphosites). Determining the characteristics of kinases that phosphorylate specific substrates have been extensively studied. Most tools utilize amino acid sequence motifs around phosphosites but do not consider the biological characteristics of substrate proteins.

**Results:** We present KSMoFinder, a kinase-substrate-motif prediction model that learns factors beyond motif similarities by integrating the biological contexts of proteins. We learn the semantics in a knowledge graph containing contextual relationships of proteins, kinase-specific motifs and motif composition, and represent the proteins and motifs as embedded vectors. Using the representations as features, we train a supervised deep learning classifier to identify kinase-phosphosite relationships. We use ground truth kinase-substrate-motif dataset from iPTMnet and PhosphositePlus and evaluate the prediction performance of KSMoFinder. Pairwise comparative assessments with prior kinase-substrate prediction tools demonstrate the superior performance of KSMoFinder. KSMoFinder trained using our knowledge graph embeddings surpasses the prediction performances using embeddings of popular protein language models such as ProtT5, ESM2 and ESM3 with a ROC-AUC of 0.851 and PR-AUC of 0.839 on a testing dataset with equal number of positives and negatives. Unlike most existing tools, KSMoFinder can be utilized to predict at the motif and at the substrate protein level.

**Availability and implementation:** All code to reproduce the results are available at https://github.com/manju-anandakrishnan/KSMoFinder. All data and KSMoFinder predictions are deposited at https://doi.org/10.5281/zenodo.15730847.

## 1 Introduction

Phosphorylation is a crucial protein post-translational modification that drives cellular processes such as cell differentiation, apoptosis, and signal transduction, wherein a phosphate molecule is transferred to specific amino acid residues of a substrate protein. Protein kinases catalyze phosphorylation and express selectivity in determining their phosphoacceptor residues [1]. Kinase dysregulation leads to abnormal substrate phosphorylation and contributes to multiple human diseases [2]. Clinical Proteomics Tumor Analysis Consortium (CPTAC) [3], PhosphositePlus (PSP) [4], and iPTMnet [5] collectively report over 300,000 unique serine, threonine, and tyrosine human phosphosites. However, less than 7% of their kinases are known [4, 5]. Therefore, uncovering kinases of human phosphosites for which kinase(s) is currently unknown is pivotal in understanding disease mechanisms.

Conventionally, popular kinase-substrate prediction tools predict kinase-specific phosphomotifs (motifs), which are short linear sequences of amino acid residues upstream and downstream of phosphosite, without considering the biological relevance between the kinase and the substrate protein [6-10]. Beyond motif specificity, phosphorylation involves either direct binding of kinases and substrate proteins or indirect interactions mediated by protein complexes [11]. Therefore relying solely on motif recognition, overlooking substrate protein’s biological characteristics may be insufficient to accurately determine kinases of phosphosites. Indeed, popular kinase-motif prediction tools such as Phosformer-ST [6] and GPS [7] suggest considering contextual factors such as protein interaction and co-expression for determining kinase specific substrates. Some recent models predict kinases at the substrate protein level without pinpointing specific phosphosites [12, 13]. However, distinct kinases often phosphorylate different phosphosites on a substrate protein. Moreover, many kinases play dual roles, as substrate proteins of other upstream kinases, which upon activation via phosphorylation regulate their downstream targets. Consequently, kinase-substrate relationship is an interplay of multiple regulating factors. Therefore, isolated predictions, at the substrate protein level or kinase-motif level may limit downstream applications such as kinase inference.

Existing kinase-substrate prediction models except Phosformer-ST generate synthetic negatives by either randomly combining kinases and substrates [8, 14] or using non-phosphorylated sites [7]. Some models randomly pair kinases of a different family and generate negatives [15], although recent studies have shown kinases of disparate families phosphorylate similar sequence motifs [9]. Random negative generation is a widely applied alternative in machine-learning tasks where ground-truth negatives are unavailable. However, in tasks such as kinase-substrate prediction, where the available knowledge is scarce (less than 7%), random generation can inject higher false negatives.

We address these challenges by developing KSMoFinder, a model for predicting kinase-phosphosite relationship by considering motif specificity at the phosphosite and substrate protein’s biological characteristics. We refer to this prediction level, as ‘substrate_motif’ level prediction as it combines features of substrate protein and motif. As an alternative to random negatives, we create biologically motivated synthetic negatives by combining two strategies, 1) pairing kinases with non-interacting substrate proteins and 2) pairing kinases with motifs it lacks specificity for. To learn the contextual relationships of proteins (kinases and substrate proteins) and motifs in a phosphorylation network, we use a knowledge graph (KG). KGs are organized graph data structures that represent information as a network of entities connected via relationships [16]. We create a phosphorylation KG by integrating diverse knowledge of kinases and substrate proteins including their function, localization and expression, and kinase-favored motifs. Using a knowledge graph embedding (KGE) algorithm, we learn the semantics in the KG and represent entities (proteins and motifs) as vectors. The representations thus generated capture the contextual links between the entities. Using these representations as features, we train a supervised classifier for identifying kinase-phosphosite relationships with ground truth positive samples from iPTMnet and PSP, and synthetically generated negatives. We evaluate KSMoFinder’s prediction performance through quantitative assessments and demonstrate its robustness and state-of-the-art (SOTA) results.

The key highlights of this work are outlined below.

i. We present a model for predicting kinases at the substrate_motif level and show its superior performance by comparing it with prior kinase-substrate prediction tools, Phosformer-ST [6], LinkPhinder [15], PredKinKG [12], and KSFinder [13].
ii. We compare our KGE with embeddings from SOTA protein language models, ESM2 [17], ESM3 [18], and ProtT5 [19] for kinase-substrate_motif prediction task and showcase the substantial prediction performance improvement using our KGE.
iii. We provide a synthetic negative dataset for kinase-substrate_motif prediction task, where the negatives are generated based on biological reasoning and data derived from experimental validation.
iv. We evaluate the influence of additional features such as kinase domain sequences and protein structures via embeddings from external models, Phosformer [14] and ProstT5 [20].
v. We show no improvement using these additional embeddings and a significant performance decline without our KGE.

We clarify the use of kinase-substrate terminologies through the rest of this paper. Phosphosite refers to a phosphorylation site on a protein. Substrate(protein) refers to a phosphorylated protein. Motif refers to the amino acid sequence around a phosphosite. Substrate_motif refers to a motif on a substrate(protein). It denotes a substrate(protein) and motif pair.

## 2 Related Work

Most existing tools learn the association between kinases and motifs without the context of the substrate(protein). Phosformer-ST [6], GPS 6.0 [7], KinasePhos 3.0 [8] are among the latest tools predicting kinases at motif level. GPS 6.0 leverages different sequence encoding methods to translate motifs to features and uses deep-learning neural network based models for predicting kinase-specific motifs [7]. KinasePhos 3.0 encodes motifs using BLOSUM62 and uses classification models to identify kinases of motifs [8]. To overcome the reliance on local sequence contexts, Phosformer-ST fine-tunes ESM2 embeddings of kinase domain sequences and phosphosite motifs using transformer architecture based model and achieves state-of-the-art performance when compared with prior kinase-motif level prediction tools [6]. The human kinase coverage of Phosformer-ST, GPS 6.0 and KinasePhos 3.0 are 300, 500 and 302 respectively. For negative generation strategies, Phosformer-ST uses experimentally determined non-favored motifs of kinases, enhancing confidence in its predictions, GPS 6.0 use non phosphorylated serine, threonine and tyrosine residues and KinasePhos 3.0 samples negatives via closed world assumption. Focussing on less studies kinases, DeepKinZero applies a zero-shot learning strategy where it transfers the knowledge learned from motif targets of 214 well-studied kinases to predict kinase-motif relationships of 112 less studied kinases [21].

Few models capture fuctional association of proteins by integrating protein-protein interaction (PPI) network along with motif sequence specificity [22, 23, 24]. PhosIDN demonstrates performance improvement when using sequence features along with PPI than using sequence features alone [22]. Phosphopredict [25] and NetKSA [26] incorportate functional features beyond protein interaction. Phosphopredict generates features by encoding motif sequence and protein functions [25]. It offers prediction coverage for 12 kinases. NetKSA combines kinase and phosphosite embeddings from two distinct networks (a phosphosite association network and a kinase association network). Its phosphosite association network is based on features such as co-occurrence, shared pathways and co-phosphorylation of the sites and the kinase association network is based on kinase interactions, common pathways and kinase family information [26]. Though NetKSA sources data for its networks based on heterogeneous relationships, its embeddings are based on homeogenous networks with presence/absence of edges between the nodes.

LinkPhinder predicts missing links in a phosphorylation network based on KG, where the nodes (kinases and substrate(protein)s) are connected by a relation (consensus motif) [15]. Using the known data of motifs, kinase family specific consensus motifs and position specific scoring matrix (PSSM) are derived using MEME (Motif-based Sequence Analysis tool). LinkPhinder provides coverage for 327 human kinases. LinkPhinder generates negatives by corrupting kinases in positive samples with a kinase from a different family. This strategy relies on the assumption that kinases of the same family phosphorylate similar motifs. While both LinkPhinder and Phosformer-ST utilize latent representations and learn features beyond local contexts, they rely solely on sequences. PredKinKG [12] and KSFinder [13] capture latent patterns in a heterogeneous phosphoproteome KG containing functional links of proteins using different graph embedding techniques and predict kinase-substrate relationships at substrate(protein) level.

KSMoFinder shares common aspects with existing tools in methodology where we utilize graph embedding for feature representation. However, it differs primarily from KSFinder in prediction level. KSFinder predicts at substrate(protein) level and includes only proteins’ function links in its KG whereas KSMoFinder predicts at substrate_motif level and includes kinase recognition motifs and amino acid residue composition of motifs in addition to proteins’ functional associations. When compared with existing tools, KSMoFinder offers the following unique features:

i. KSMoFinder combines functional features of kinases and substrate(protein)s in addition to motif sequence features and predicts kinases at the substrate_motif level. Although LinkPhinder and PhosIDN predict at the substrate_motif level, neither considers different biological associations of proteins. NetKSA learns functional associations of phosphosites rather than relationships between kinase and substrate(protein).
ii. KSMoFinder provides an broad kinase coverage including 430 human kinases across 9 kinase groups, Atypical, AGC, CAMK, CK1, CMGC, STE, TKL, TK and Other.
iii. Our negative generation strategy is motivated by compelling biological rationale and data based on experiments. For kinase-motif negative samples, we utilize experimentally derived unfavored motifs of kinases. For kinase-substrate(protein) negatives, we pair non-interacting proteins. Moreover, our negatives include data where a positive kinase-motif is paired with non-interacting substrate(protein)s and a positive kinase-substrate(protein) is paired with unfavored motifs.

To the best of our knowledge, KSMoFinder is the first model to offer substrate_motif level prediction with broad human kinome coverage, and to base its predictions on both substrate(protein)’s characteristics and motif sequence.

## 3 Methods

### Development of KSMoFinder

KSMoFinder comprises of two major components – 1) a knowledge graph embedding model, which learns the semantics of a phosphorylation network and embeds the entities in a latent vector space; 2) a supervised neural network classifier, which learns from the embedded vectors of kinases, substrate(protein)s and motifs and predicts phosphorylation probability (Figure 1).

**Figure 1.**
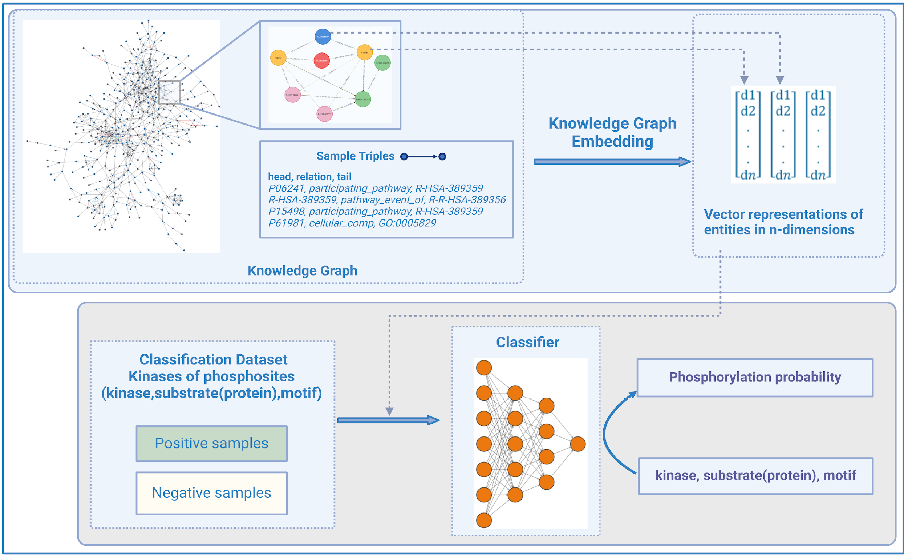
Overview of KSMoFinder: a kinase-phosphosite prediction model consisting of a knowledge graph embedding and a neural network classifier.

#### 3.1.1 Knowledge graph

Our knowledge graph, a semantic network of kinases, substrate(protein)s and motifs is created by integrating relevant contextual information of proteins and motifs from different sources. It comprises 4,881,408 unique triples, 32 relationships, and 360,098 nodes. Each triple is modeled to present a semantic meaning using a head entity (h), a relation (r) and a tail entity (t). For example, the participation of protein, ‘14-3-3 protein gamma’ in the pathway, ‘activation of BAD and translocation to mitochondria’ is represented using the triple, ‘P61981|participating_pathway|R-HSA-111447’, where P61981 and R-HSA-111447 are identifiers of the involved protein and pathway respectively. Table 1 details the different relationships in our KG with a description, triple count for each relationship and an example triple.

**Table 1.** Lists of the relationships in the KSMoFinder knowledge graph, along with example triples, and triple counts for each relationship.

| Relationship | Relationship description | Example Triple | Triple Description | Triple count |
| --- | --- | --- | --- | --- |
| participating_pathway | Pathways, a protein participates in | P61981\|participating_pathway\|<br>R-HSA-111447 | 14-3-3 protein gamma participates in the pathway, Activation of BAD and translocation to mitochondria. | 55,637 |
| part_of_complex | Protein and protein complexes it is associated with | P08603\|part_of_complex\|<br>R-HSA-1006173 | Complement factor H protein is part of the complex, Host cell surface [plasma membrane]. | 89,165 |
| pathway_event_of | Pathway hierarchies, where one pathway is part of another pathway | R-HSA-73943\|pathway_event_of\|<br>R-HSA-73942 | Reversal of alkylation damage by DNA dioxygenases is a subclass of DNA damage reversal. | 2,574 |
| bio_process | Processes, a protein carries out | O75342\|bio_process\|GO:0061436 | Arachidonate 12-lipoxygenase, 12R-type is involved in the process, establishment of skin barrier. | 138,446 |
| regulates | A biological process regulates another biological process | GO:0035542\|regulates\|<br>GO:0035493 | SNARE complex assemble is regulated by a biological process, regulation of SNARE complex assembly. | 2,298 |
| positively_regulates | A biological process regulates another biological process, positively | GO:0060045\|positively_regulates\|<br>GO:0060038 | Cardiac muscle cell proliferation is regulated by positive regulation of cardiac muscle cell proliferation. | 1,811 |
| negatively_regulates | A biological process regulates another biological process, negatively | GO:0016248\|negatively_regulates\|<br>GO:0015267 | Channel activity is regulated by channel inhibitor activity. | 1,587 |
| mol_func | Functions, a protein is involved in | Q9UMQ6\|mol_func\|GO:0005509 | Calpain-11 is involved in calcium ion binding. | 57,910 |
| capable_of | Inferred relationship where a protein complex can be involved in another function/process | GO:0106068\|capable_of\|<br>GO:0019789 | SUMO ligase complex is capable of being involved in SUMO transferase activit. | 265 |
| capable_of_part_of | Inferred relationship where a protein complex can be part of another function/process/cellular component | GO:0005840\|capable_of_part_of\|<br>GO:0006412 | Ribosomes are capable of being a part of protein translation. | 169 |
| cellular_comp | Cellular components, a protein resides in | P61981\|cellular_comp\|<br>GO:0005737 | 14-3-3 protein gamma is located in cytoplasm. | 81,635 |
| occurs_in | Cellular component, a function/process occurs in | GO:0030527\|occurs_in\|<br>GO:0000785 | Functions contributing to the chromatin's structure occur in chromatin. | 194 |
| part_of | Function/process is part of a biological process (or) cellular component is part of another cellular component | GO:0046943\|part_of\|<br>GO:1905039 | carboxylic acid transmembrane transporter activity is part of carboxylic acid transmembrane transport. | 4,427 |
| expressed_in | Tissues in which a protein is expressed in | Q9UFE4\|expressed_in\|<br>nasopharynxrespiratory-epithelial-cells | CCDC39 is expressed in nasopharynxrespiratory-epithelial-cells. | 85,557 |
| is_a | A cellular component/process/function has a transitive relationship with another component/process/function | GO:0051006\|is_a\| GO:0051004 | positive regulation of lipoprotein lipase activity is a regulation of lipoprotein lipase activity. | 35,020 |
| has_domain | Domains, a kinase is associated with | P54646\|has_domain\|IPR032270 | PRKAA2 has the domain AMPK, C-terminal adenylate sensor domain. | 1,622 |
| belongs_to_family | Kinase families, a kinase belongs to | Q05513\|belongs_to_family\|PKC | PRKCZ belongs to the protein kinase C family. | 406 |
| homologous_superfamily | Homologous superfamilies (usually shared tertiary structures), a kinase belongs to | Q13177\|homologous_superfamily\|<br>IPR011009 | Serine/threonine-protein kinase PAK 2 belongs to the homologous superfamily, protein kinase-like domain superfamily. | 993 |
| is_a(domain) | A domain is a sub-class of another domain in the hierarchy | IPR035851\|is_a(domain)\|IPR000980 | Tyrosine-protein kinase HCK, SH2 domain is a SH2 domain. | 171 |
| is_a(family) | A kinase family is a sub-class of another family in the hierarchy | Trbl\|is_a(family)\|CAMK | Tribbles subfamily is a CAMK Serine threonine kinase family. | 136 |
| is_a(form) | Canonical protein form of an isoform/a proteoform | Q8NHQ8-2\|is_a(form)\|Q8NHQ8 | Ras association domain-containing protein 8 (isoform 2) is a form of the protein ras association domain-containing protein 8. | 8,164 |
| k_specific_motif | 9-mer motif, a kinase is known to have specificity to | Q9BWU1\|k_specific_motif\|<br>LDLFSPSVT | Cyclin dependent kinase, CDK19 has phosphorylation specificity to the 9-mer motif, LDLFSPSVT. | 1,329,571 |
| has_motif | Substrate(protein), the 9-mer motif is on | Q9Y4B6\|has_motif\|YDIQTGNKL | DDB1- and CUL4-associated factor 1 has a 9-mer motif, YDIQTGNKL. | 347,109 |
| residue_1... residue_4 | Amino acid residues upstream of the phosphosite | SSPSTPVGS\|residue_1\|aa_S | Amino acid at four residues upstream of the phosphosite is Serine. | 1,171,796 |
| residue_5 | Amino acid, at the phosphosite | SSPSTPVGS\|residue_5\|aa_T | Amino acid at the phosphosite is Threonine. | 292,949 |
| residue_6.. residue_9 | Amino acid residues downstream of the phosphosite | SSPSTPVGS\|residue_6\|aa_P | Amino acid at one residue downstream of the phosphosite is Proline. | 1,171,796 |

The two primary data types of our KG are kinases and phosphosites. Annotated human kinases and phosphosites are retrieved from PSP [4] and iPTMnet [5]. In addition, a large number of experimentally determined human phosphosites from different CPTAC Proteomics Data Commons studies are extracted [3]. Motifs of length 9 (9-mer) are constructed using four residues upstream (-4) and downstream (+4) of phosphosites. Independent of phosphosite data collection, from two in-vitro experimental studies that profiled substrate specificities of kinases using synthetic peptide arrays, kinase specific motifs are obtained [9, 10]. When the kinase/substrate(protein) is an isoform, the canonical form of the protein is obtained from UniProt [27]. The proteins’ functions are obtained from 1) molecular functions (Gene Ontology), biological processes (Gene Ontology), participating pathways (Reactome) from UniProt [27]; and participating protein complexes from Reactome [28]. The proteins’ localization and expression data are sourced from 1) cellular location from UniProt [21]; and 2) tissue of expression from Human Protein Atlas [29]. Kinase domain data (domain names) and homologous superfamily information are obtained from InterPro [30]. We use kinase family information in Phosformer [14]. The hierarchical information on pathways, protein complexes are retrieved from Reactome [28]; biological processes, molecular function, cellular components are retrieved from Gene Ontology [31, 32]; kinase domains and kinase homologous superfamilies are obtained from InterPro [30]. The 9-mer motifs are represented by linking them to constituting amino acids at each position. For example, motif, ‘SSPSTPVGS’ connects with 9 amino acid nodes representing serine at positions 1, 2, 4, and 9; proline at positions 3 and 6; threonine at 5; valine at 7; and glycine at 8. Supplementary File 1 details the steps involved in creating our KG.

#### 3.1.2 Knowledge graph embedding

We leverage Pykeen library [33] and develop four independent KGE models using different algorithms: 1) TransE [34], 2) DistMult [35], 3) ComplEx [36], and 4) ExpressivE [37], and learn the semantics in the KG. Each algorithm uses different scoring functions to achieve an optimal representation of the network entities. TransE forces the sum of the head and relation vectors to equate to the tail vector using the L1 norm [34]. DistMult and ComplEx are bilinear embedding models that represent entities using matrix multiplication. DistMult scores triple as the product of head, relation, and tail vectors, which yields the same score when head and tail entities swap positions [35]. ComplEx algorithm differs from DistMult by embedding in complex number space where the entity’s vector is used when in the head position and its conjugate is used when in the tail position [36].

ExpressivE, a recently published KGE algorithm, represents relation as hyper-parallelogram and head and tail entities as vectors respectively and captures complex relationships such as hierarchy in relations [37].

We split the KG into training and validation datasets. A subset of relationship types that directly impact kinases, substrate(protein)s, and motifs are included in the validation dataset. The relationship types used for KG validation are provided in Supplementary File 1. Due to the large graph size, 2.5% of the triples of each relationship type are randomly sampled and a validation dataset is created with 11,186 triples. The KGE models are trained with 4,870,222 triples using the hyperparameters (embedding dimension of 100; batch size of 25,000; Adam optimizer; PairWiseHinge loss; 1 negative triple per positive triple; learning rate of 1e-03). The optimal number of training iterations is determined by evaluating the validation dataset for an increase in mean reciprocal rank (MRR) for every 20 iterations with a patience count of 2. During evaluation, each positive triple in the validation dataset is individually ranked with respect to the negative triples generated by corruption of positive entities. By ranking the correct entity among all the evaluated entities, a validation triple’s reciprocal rank is computed. The average reciprocal rank of all validation triples is MRR.

#### 3.1.3 KSMoFinder classification dataset

We generate a classification dataset with positive and negative samples of kinase-substrate_motif phosphorylation events (Figure 2). The positive dataset is obtained from PSP [4] and iPTMnet [5]. PSP’s kinase-phosphosite dataset contains experimentally determined substrates and cognate kinases curated from literature [4]. iPTMnet is an integrated resource for post-translational modifications and includes kinase-substrate relationships from multiple sources such as Phospho.ELM, neXtProt, UniProt, Signor, HPRD, PhosphoGRID [5]. From iPTMnet, we retrieve kinase-phosphosite data that have a confidence score >= 1. 9-mer motifs are constructed using four residues upstream and four residues downstream of phosphosites. Each classification data sample is represented by <k, s, m, l>, where ‘k’ denotes the kinase, ‘s’ the substrate(protein), ‘m’ the motif, and ‘l’ the label. Label is ‘1’ for positive and ‘0’ for negative.

**Figure 2.**
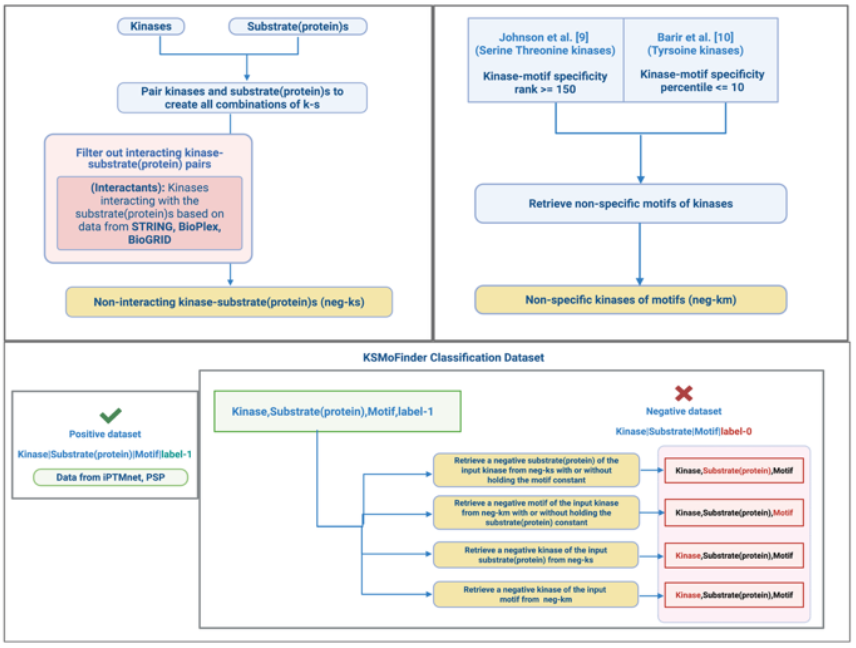
KSMoFinder classification dataset creation workflow: (a) construction of neg-ks. (b) creation of neg-km, (c) positive dataset sourced from iPTMnet and PSP and negative dataset generated by sampling data from neg-ks and neg-km.

We create a negative dataset by combining the following two strategies.

i. pairing kinase and substrate(protein) with no evidence of interaction based on data from STRING [38], BioPlex [39] and BioGRID [40]. This dataset is referred to as ‘negative-kinase-substrate (neg-ks)’ (Figure 2.a). When extracting interactants from STRING, only those protein pairs with a confidence score >= 0.75 are considered to reduce false positive interactants. Supplementary File 1 details the steps involved in extracting protein interactants.
ii. pairing kinases with motifs that show no phosphorylation specificity based on datasets published by Johnson et al. [9] for serine-threonine kinases and Barir et al. [10] for tyrosine kinases. From serine/threonine kinases dataset kinase-motif specificity with a rank of 150 or greater are selected as negative kinase-motif pairs. For tyrosine kinases, kinases with a percentile <=10 for a peptide are selected as negative kinase-motif pairs. This dataset is referred to as ‘negative-kinase-motif (neg-km)’ (Figure 2.b).

Based on the three entities (kinase, substrate(protein) and motif) in each positive sample, negatives are generated using the following methods (Figure 2.c.)

i. For the motif in the positive sample, a random kinase is selected from ‘neg-km’.
ii. For the substrate(protein) in the positive sample, a random kinase is selected from ‘neg-ks’.
iii. For the kinase and substrate(protein) combination in the positive sample, a random motif is selected from ‘neg-km’.
iv. For the kinase and motif combination in the positive sample, random substrate(protein) is selected from ‘neg-ks’.

When no data is available for (iii) or (iv), a random sample with negative motif or negative substrate(protein) of the kinase is chosen. In all cases, we ensure the substrate_motif pair is a valid phosphosite. This results in 19,574 positive and 69,934 negative samples. For each kinase, we randomly sample 20% of its positives and negatives for the testing dataset. The resulting training dataset consisted of 16,137 positives and 56,068 negatives. We create two testing datasets, 1) with 1:1 ratio of positives to negatives (Testing Dataset 1) and 2) with the distribution of positives to negatives similar to the training dataset (Testing Dataset 2).

Our KG does not include direct links between kinase and substrate(protein). The kinase-motif triples in the KG exclude instances that are part of the positive classification dataset. This is a measure to avoid data leakage in our embeddings and thereby to KSMoFinder classifier.

#### 3.1.4 KSMoFinder Classifier

Using embeddings from KGE as features, a neural network model is trained to predict the kinases of substrate_motif (Figure 3). The vector representation of the sample *<k, s, m>* is denoted as *<V*_*k*_, *V*_*s*_, *V*_*m*_>. Two bilinear transformation layers individually capture the associations between kinase and substrate(protein), and kinase and motif. The two individual combinations with kinases are represented as,

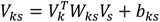

where, V_ks_ is the resultant vector combining the kinase vector and the substrate(protein) vector, W_ks_ is the associated learned weight matrix and b_ks_ is the bias.

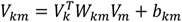

where, V_km_ is the resultant vector combining the kinase vector and the motif vector, W_km_ is the associated learned weight matrix and b_km_ is the bias.

**Figure 3.**
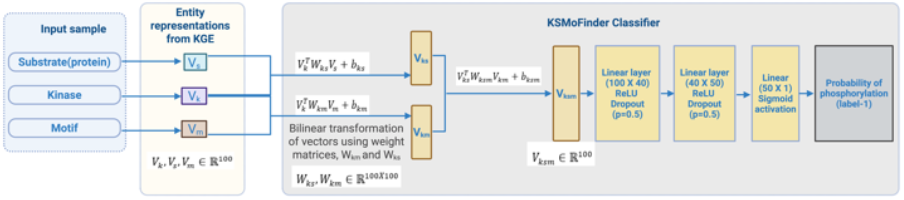
KSMoFinder classifier architecture, where kinase vector is combined with motif vector and substrate(protein) vector, transformed and passed further through neural network layers.

V_ks_ and V_km_ are then transformed with a third bilinear layer resulting in the vector, V_ksm_, which is passed through two linear hidden layers of 40 and 50 neurons, each followed by a rectified linear unit, non-linear activation function and a drop-out layer with a probability of 0.5 for training regularization. The output is transformed using the sigmoid activation function to compute the probability for positive class.

The loss between the predicted and actual output is computed using binary cross entropy (BCE).

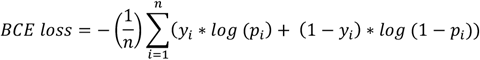

where, p_i_ is the predicted probability of positive class, y_i_ is the true class label, and n is the number of samples.

### Performance Evaluations

#### 3.2.1 Evaluation of different knowledge graph embeddings

Two assessments are conducted to assess the best feature representations. Assessment 1 evaluates embeddings of the four KGE models (TransE, DistMult, ComplEx, ExpressivE) trained on our KG. We use KSMoFinder classification dataset for training and assessment. The training dataset is of size 72,205 (positives: 16,137; negative: 56,068). Testing dataset 1 consists of 6,874 samples (positives: 3,437; negatives: 3,437), and Testing dataset 2 consists of 16,559 samples (positives: 3,437; negatives: 13,866). The KGE model with the best performance in this assessment is selected as the optimal embedding for KSMoFinder and is referred to as ‘KSMoFinder-KGE’.

Assessment 2 compares KSMoFinder-KGE with embeddings from ESM2 [17], ESM3 [18] and ProtT5 [19]. ProtT5-XL-UniRef50 (ProtT5) and ESM2 are popular protein embedding models trained on a large corpus of protein sequences [19]. ESM2 uses masked prediction tasks for learning sequence contexts and represents evolutionarily conserved functions and proteins’ structures in its embeddings [17]. ESM3 is a recent generative model combining structure, function, and sequence modalities in its protein representation. It learns each modality from discrete encoded tokens such as atomic structures, functional keywords (domains, enzymatic activity), and protein sequences [18]. The protein sequences of kinases, substrate(protein)s and motifs are obtained from UniProt. Using the sequences as input, embeddings are retrieved from ESM2 and ESM3. To fetch embeddings of kinases and substrate(protein)s from ProtT5, their UniProt identifiers are passed as input. For motifs, residue level embedding is retrieved from ProtT5. Each residue level embedding yields a vector of length 1024, so the mean of residue-level embeddings of the 9-mer motif is computed. As Assessment 2 investigates embeddings from multiple external sources, only those samples with embeddings from all the sources are included in training and testing datasets for this assessment. This results in a training dataset of size 48,349 (positives: 14,019; negative: 34,330). Testing dataset 1 and Testing dataset 2 are of sizes 6,050 samples (positives: 3,025; negatives: 3,025) and 11,514 samples (positives: 3,025; negatives: 8,489) respectively. To rule out inherent bias in the classification dataset, we generate random embeddings of size 100 for each entity (kinases, substrate(protein)s and motifs), train a model and evaluate its performance.

#### 3.2.2 Evaluation with embeddings of additional features

We assess the inclusion of additional features via embeddings that might aid the model in learning patterns about kinases phosphorylating substrate_motif. Different evaluations are conducted using kinase domain sequences, protein structures, and 15-mer motifs (instead of the 9-mer motifs used in the original KGE for direct comparison with Phosformer). These evaluations are grouped under ‘Assessment 3’.

To assess the contribution of kinase domain sequences, we incorporate embeddings from Phosformer [14], a transformer architecture-based model pre-trained on kinase domain sequences and substrate(protein) sequences. The kinase domain sequences’ embeddings and motif embeddings are extracted from Phosformer’s pre-trained model [14], combined bilinearly with KSMoFinder-KGE after scaling their features to KSMoFinder-KGE’s embedding range, and a classifier model is trained.

For structure information, we use ProstT5, a bilingual protein language model trained to translate protein sequences to structures, and vice versa, which is used to learn the structure of kinases and substrate(protein)s [19]. To translate from the sequence to structure, protein sequences are prefixed with ‘<AA2fold>‘ and passed to ProstT5. As we are interested in the representation rather than the predicted structure, embeddings from ProstT5’s last layer are retrieved. After feature scaling, ProstT5 embeddings are combined with KGE embeddings via bilinear transformation, and the output is fed through subsequent neural network layers. We combine embeddings from all three pre-trained models (KSMoFinder-KGE, ProstT5, and Phosformer), and evaluate them for performance improvement. Finally, we assess 1) the effect of dropping KSMoFinder-KGE by using ProstT5 and Phosphormer representations alone and 2) contribution of biological associations by dropping protein representations from KSMoFinder-KGE.

The training dataset used for this assessment is of size 65,962 with 15,565 positives and 50,397 negatives. Testing dataset 1 consists of 6,680 samples (positives: 3,340; negatives: 3,340), and Testing dataset 2 consists of 15,447 samples (positives: 3,340; negatives: 12,107).

#### 3.2.3 Comparative evaluation of KSMoFinder with other kinase-substrate prediction models

We evaluate KSMoFinder at the motif level by comparing its performance with Phosformer-ST, which has demonstrated superior performance over prior kinase-motif level prediction tools [6]. As KSMoFinder predicts at the substrate_motif level, our negative dataset contains samples where the kinase is paired with a non-interacting substrate(protein), although the kinase-motif combination is true. For a fair comparison, we eliminate such samples from the negative dataset and retain only those samples with unfavored motifs of kinases. LinkPhinder uses a knowledge graph model and predicts novel links between kinases and phosphosites [18]. We compare KSMoFinder with LinkPhinder at the substrate_motif level and with PredKinKG [12] and KSFinder [13] at the substrate(protein) level.

To facilitate comparisons with prediction models at substrate(protein) level, from the negative dataset, we remove samples where the kinase-substrate(protein) combination is true. Our test dataset may have samples that the other models are trained on, giving an advantage to those models over KSMoFinder. These evaluations are grouped under ‘Assessment 4’.

#### 3.2.4 Comparative evaluations without easy test scenarios

Kinases with same domains or belonging to the same family are often known to recognize similar motifs. Although our KG has no knowledge of kinase sequences, it has links between kinases and their domains, and kinases and their families. We refer to kinases belonging to the same family or containing the same domain as ‘similar kinases’. Test scenarios where a motif is paired with a similar kinase and label as in the training sample are referred to as ‘easy test scenarios’. For example, given *k*_*1*_ and *k*_*2*_ are similar kinases, sample, *<k*_*1*_, *s, m, l>* in test dataset is an easy test, if *<k*_*2*_, *m>* exists in the training dataset only in the class label, *l*. If *<k*_*2*_, *m>* exists in both class labels, 0 and 1 in the training dataset, then it is not considered an easy test. We conduct tests without easy test scenarios for Assessment 2 and Assessment 4 where we compare performance of our embedding and model with others.

For all comparisons, we bootstrap 1000 samples with replacement from the testing datasets to calculate confidence intervals of ROC-AUC and PR-AUC scores.

## 4 Results

### 4.1 Comparative performance evaluation of KGE models (Assessment 1)

The KGE models are trained individually on an NVIDIA A100 GPU with 40 GB memory. A constant MRR score with increasing training iterations indicates the optimal epoch. We evaluate the best KGE by assessing it on our downstream task of predicting kinase-substrate_motif relationships. Based on the evaluation results summarized in Table 2, representations embedded by the TransE algorithm are selected as optimal for KSMoFinder.

**Table 2.** Comparative evaluation of embeddings from four KGE models in identifying kinase-phosphosite relationship.

| Algorithm | Testing dataset 1 |  | Testing dataset 2 |  |
| --- | --- | --- | --- | --- |
|  | ROC-AUC | PR-AUC | ROC-AUC | PR-AUC |
| TransE | **0.905±0.0** | **0.903±0.001** | **0.922±0.00** | **0.78±0.001** |
| DistMult | 0.871±0.001 | 0.864±0.001 | 0.896±0.0 | 0.683±0.001 |
| CompLex | 0.79±0.001 | 0.741±0.001 | 0.814±0.0 | 0.514±0.01 |
| ExpressivE | 0.555±0.001 | 0.528±0.001 | 0.561±0.001 | 0.221±0.0 |

### 4.2 Comparative performance evaluation of KSMoFinder-KGE with other embeddings (Assessment 2)

Table 3 summarizes the results of Assessment 2 where we compare KSMoFinder’s embeddings with embeddings from 1) ProtT5, 2) ESM2, 3) ESM3, and 4) Random generation. The model KSMoFinder developed using KSMoFinder-KGE embeddings outperformed the models developed using embeddings from ProtT5, ESM2, and ESM3. The scores approximating 0.5 on the randomly generated embeddings and classification testing dataset 1, which contains 1:1 ratio of positives to negatives, demonstrates the unbiased nature of our classification dataset.

**Table 3.** Comparing the embeddings from KSMoFinder KGE with other protein-based models in identifying kinase-phosphosite relationship.

| Embedding source | Testing dataset 1 |  | Testing dataset 2 |  |
| --- | --- | --- | --- | --- |
|  | ROC-AUC | PR-AUC | ROC-AUC | PR-AUC |
| KSMoFinder-KGE | **0.851±0.008** | **0.839±0.008** | **0.914±0.003** | **0.79±0.009** |
| ProtT5 | 0.752±0.007 | 0.726±0.001 | 0.871±0.04 | 0.676±0.009 |
| ESM2 | 0.691±0.009 | 0.659±0.01 | 0.854±0.003 | 0.615±0.008 |
| ESM3 | 0.501±0.003 | 0.5±0.002 | 0.493±0.002 | 0.26±0.001 |
| Random | 0.498±0.004 | 0.5±0.002 | 0.497±0.003 | 0.263±0.001 |

Phosphorylation is often a part of the cascade of signaling events and is influenced by regulating factors. Unlike ESM2 [17] and ProtT5 [19], which primarily rely on protein sequences, our approach extracts features by learning implicit semantics in a KG focused on proteins’ biological connections, enriching the contextual information of proteins in KSMoFinder-KGE. ESM3 [18] addresses this limitation by training on functional annotations of proteins and residues, however, our assessment of ESM3 for kinase-substrate_motif prediction shows poor performance. ESM3 is trained using abstract keywords (binding, catalytic sites, post-translational modifications), which may not be sufficient for specific tasks such predicting kinases of phosphosites.

Furthermore, KSMoFinder-KGE is of smaller dimension with a size of 100 whereas ProtT5, ESM2, and ESM3 are 1024, 1280, and 1536 respectively. Despite the smaller embedding size, which lowers computation resources, KSMoFinder-KGE shows superior performance.

### 4.3 Evaluation with embeddings of additional features (Asessment 3)

Phosformer and Phosformer-ST utilized kinase domain sequences to predict the kinase-motif relationship [6, 14]. The classifier developed using kinase domain sequences’ embeddings and 15-mer motif from Phosformer’s pre-trained model doesn’t improve performance further (Supplementary File 2 (Table 1) - Test No.1 vs Test No.3). Although KSMoFinder-KGE has no direct knowledge of kinase domain sequences, it is trained on kinase domain names and domain hierarchies. Presumably, Phosformer’s embeddings on kinase domain sequences do not offer more meaningful features than what has already been learned by KSMoFinder-KGE.

Leveraging the pre-trained model, ProstT5, which learned AlphaFold2 predicted 3D protein structures, the effect of including structural features of kinases and substrate(protein)s is evaluated (Supplementary File 2 (Table 1) - Test No.1 vs Test No.2). There is no performance increase in models trained with kinase domain sequences or structural features, suggesting KSMoFinder-KGE is sufficient. Next, to investigate whether embeddings from Phosformer and ProstT5 are adequate on their own, a model is trained without KSMoFinder-KGE. This investigation is necessary to assess the contribution from KSMoFinder-KGE. The embeddings from the two models: 1) representations of kinases and substrate(protein)s from ProstT5, and 2) representations of 15-mer motifs and kinase domain sequences from Phosformer are utilized. The substantial drop in prediction performance in this assessment demonstrates the enriched feature contribution by our KGE (Supplementary File 2 (Table 1) - Test No.4 vs Test No.5). Using Testing dataset 1, ROC-AUC drops from 0.893 to 0.724 and PR-AUC drops from 0.886 to 0.652. Using Testing dataset 2, ROC-AUC drops from 0.92 to 0.802 and PR-AUC drops from 0.777 to 0.422. We also observe a reasonable performance decrease when excluding learned representations of proteins’ biological associations (Supplementary File 2 (Table 1) - Test No.2 vs Test No.6).

The optimal hyperparameters used to develop the different classifier models for all the assessments are provided in Supplementary File 3.

### 4.4 Comparative Evaluation of KSMoFinder with other ki-nase-substrate prediction tools (Assessment 4)

We benchmark KSMoFinder against four other kinase-substrate prediction tools, LinkPhinder, Phosformer-ST, KSFinder, and PredKinKG. Due to the lack of benchmarking datasets, KSMoFinder classification dataset is used for comparison. The substrate(protein) and motif specifics in our dataset allow comparative evaluation of tools that predict kinase-substrate relationship at the protein level, the motif level, or both. For motif level evaluation, we restrict our comparison to Phosformer-ST, as it showed superior performance over other kinase-motif level prediction tools. A direct comparison of the different tools is impossible, as the tools differ in prediction levels and utilize different datasets.

To facilitate a fair comparison with KSMoFinder, we performed several pre-processing strategies. These include 1) eliminating negatives where either kinase-substrate(protein) or kinase-motif is a positive sample, 2) not excluding other models’ training samples from the testing dataset, and 3) limiting to the subsets of data that the respective models are trained on, whenever possible. KSMoFinder is compared pairwise with the other models. The evaluation results using a testing dataset containing a 1:1 ratio of positives to negatives are reported in Figure 4.

**Figure 4.**
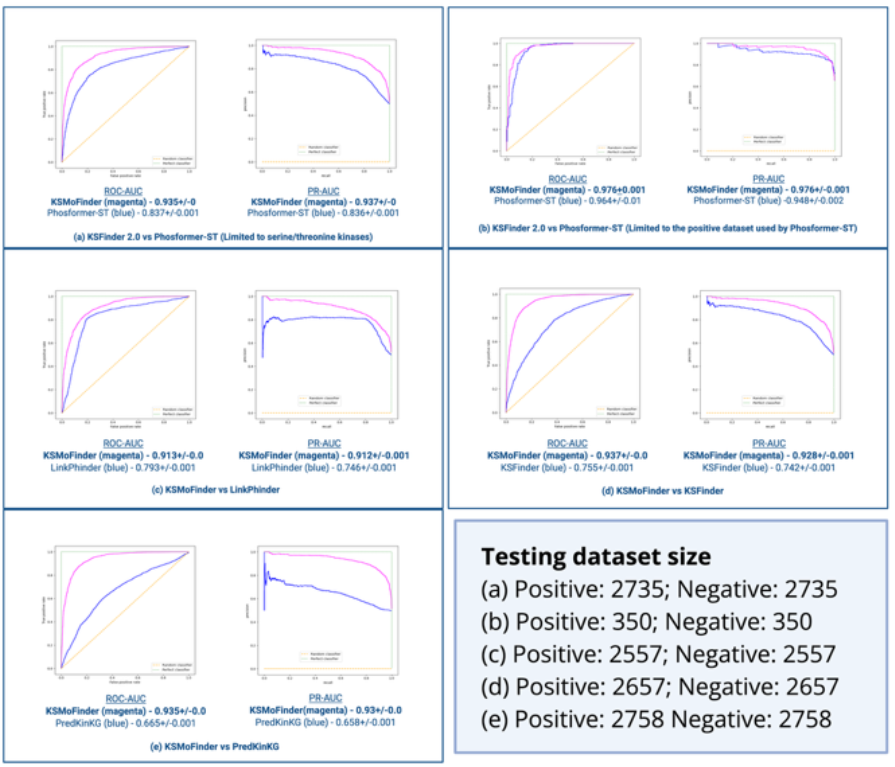
Pairwise comparative evaluation results of KSMoFinder with other models using Testing Dataset 1. Scores are reported in ROC-AUC and PR-AUC. (a) KSMoFinder vs Phosformer-ST using the positive dataset from iPTMnet and PSP, (b) KSMoFinder vs Phosformer-ST using the subset containing positive samples reported by Johnson et. al., [4], (c) KSMoFinder vs LinkPhinder, (d) KSMoFinder vs KSFinder, (e) KSMoFinder vs PredKinKG.

Our comparison with Phosformer-ST is limited to the 300 serine-threonine kinases it is trained on. Phosformer-ST uses experimentally profiled kinase-specific motifs reported by Johnson et al [6]. We perform two sub-assessments to compare KSMoFinder with Phosformer-ST. In sub-assessment 1, positive kinase-motif pairs are sourced from iPTMnet and PSP, and KSMoFinder outperforms Phosformer-ST (Figure 4.a). Sub-assessment 2 is limited to positive kinase-motif pairs reported by the work of Johnson et al., the dataset Phosformer-ST is trained on. KSMoFinder shows similar performance as Phosformer-ST in sub-assessment 2 (Figure 4.b). This demonstrates KSMoFinder’s generalization and exceptional prediction capability compared to the SOTA kinase-motif prediction model. KSMoFinder also shows superior performance over LinkPhinder (Figure 4.c) and prior protein-level prediction models, KSFinder and PredKinKG (Figures 4.d, 4.e). We also evaluate the models using testing datasets with higher negatives than positives. The results of this evaluation are provided in Supplementary File 2 (Figure 1). The relative performance of KSMoFinder in comparison with other tools is similar to the results with a 1:1 ratio of positives to negatives.

Prediction performance of KSMoFinder on different kinase groups, families and individual kinases are reported in Supplementary File 4. The results of evaluations with and without easy test scenarios are reported in Supplementary File 2 (Tables 2, and 3). As expected, there was a decrease in performance in tests without easy test scenarios for most models except LinkPhinder. However, there is no effect on the relative performance of KSMoFinder in comparison with models.

### 4.5 Analysis of KSMoFinder predictions

As KSMoFinder learns biological links of kinases and substrate(proteins)s such as pathway participation, co-location, co-expression, common processes and functions, its predictions for the same kinase-motif pair varies depending on the substrate(protein)’s context. For example, the 9-mer motifs surrounding the phosphosites, S211 on GULP1, ‘MTPKSPSTD’, S215 on PROSER2, ‘LSPTSPFRE’, and S75 on CRYBG1, ‘ASAASPESK’ are favored motifs of CDK19 based on synthetic peptide profiling [9], but the proteins have no known functional association with CDK19. KSMoFinder predicts phosphorylation of these sites by CDK19 with low probabilities of 0.465, 0.209 and 0.329. Supporting these predictions, inhibition of CDK19 and CDK8 in HCT116 cells shows no difference in phosphorylation abundance at S211 on GULP1, S215 on PROSER2 and S75 on CRYBG1 [41]. However, the same experiment shows more than two-fold decrease in phosphorylation at S32 on AFF4, S314 on MED26, S1112 on MED14 and KSMoFinder predicts CDK19 phosphorylation of these sites with a probability > 0.9. All three proteins and CDK19 are located in the nucleus. MED14 and MED26 participate with CDK19 in common pathways such as transcription regulation of adipocyte differentiation and RSV-host interactions.

Other sites, including S481 on MED13, S82 on MED8, S505 on MELK have motifs not favored by CDK19 [9], but the proteins have no obvious biological links with CDK19. As expected, these sites which show no difference in phosphorylation abundance in the perturbed experiment are predicted with lower probabilities of phosphorylation by CDK19.

Similarly, Leucine-rich repeat serine/threonine-protein kinase 2 (LRRK2) is predicted to phosphorylate motif, ‘RTPRTPRTP’ on la-related protein 1 (LARP1) with a high probability of 0.863. Whereas the predicted probability of the same motif on MED13 by LRRK2 is 0.002. Correlating with these predictions, there are no obvious biological links between MED13 and LRRK2 whereas LARP1 and LRRK2 are involved in common biological processes such as macroautophagy, cellular response to starvation and cell proliferation.

## 5 Discussion

In this work, we present KSMoFinder, a novel model for predicting kinase-phosphosite relationship. Unlike most prior kinase-substrate prediction models, which considers either motif or substrate(protein) features, KSMoFinder combines features of substrate(protein) and motif and offers predictions at substrate_motif level. KSMoFinder offers prediction coverage for 430 human kinases spanning across 9 kinase groups. This presents a major advancement for kinase-substrate predictions as KSMoFinder is the first model to offer this broad kinome coverage with substrate_motif level prediction. Based on the collective data from iPTMnet and PSP, about 27% of the phosphosites are phosphorylated by more than one distinct kinase. By modeling the task as a binary classification, our model learns about phosphorylation sites targeted by multiple kinases.

Comparative assessment of our KGE with embeddings from sequence based protein models, ESM2 and ProtT5 underscore the importance of integrating context-specific functional features for kinase-substrate prediction tasks. KSMoFinder’s negatives are created based on a biological rationale and includes samples simulating realistic patterns in kinase-phosphosite relationships. KSMoFinder’s ability to make predictions at substrate_motif level allowed comparison with other models offering predictions at the motif, the substrate and the substrate_motif levels. These other models, LinkPhinder and Phosformer-ST use protein sequence representations and KSFinder and PredKinKG use functional features for predicting kinases of substrates. KSMoFinder’s superior performance over these models further demonstrates the strength of combining protein functions and motif sequence features in its representations.

Including structural features did not improve KSMoFinder’s performance further. One possible explanation for this finding is that phosphosites are located predominantly in intrinsically disordered regions of proteins lacking stable structures [42] whereas ProstT5 is trained on stable structures. Advanced structural embeddings capturing the dynamic structural conformation of proteins may be necessary for learning the structural factors affecting phosphorylation. Further, our KG contains relationships linking kinases and their homologous superfamily, which allows learning implicit similarities in tertiary structures of kinases.

## 6 Conclusions

Our study addresses a major limitation in kinase-substrate prediction by incorporating factors beyond motif sequence specificity. KSMoFinder’s broad human kinome coverage and its ability to predict at the substrate_motif level shows promise for its application in phosphorylation research. Applications such as kinase inference relying on kinase-substrate association could benefit from KSMoFinder’s predictions which are based on proteins’ contextual relationships in addition to motif specificity.

