## Supplementary File 1 for "KSMoFinder - Knowledge graph embedding of proteins and motifs for predicting kinases of human phosphosites"

**KSMoFinder knowledge graph creation steps**

Our knowledge graph is created with data from different databases and scientific literature. Table 1 presents the high-level steps involved in data collection and data source. Following Table 1, we provide the detailed steps and data download/retrieval date from the source.

Table 1. Lists the steps involved in collecting the data for constructing the knowledge graph. It presents the types of data and the corresponding data source.

| **Steps** | **Data** | **Data Source** |
| --- | --- | --- |
| Step 1. Collect, kinases, substrate proteins and phosphorylation sites | Phosphosites (substrate(protein), site of phosphorylation) | iPTMnet [1], PSP [2], CPTAC-PDC [3] |
|  | Kinases | iPTMnet [1], PSP [2] |
| Step 2. Convert phosphorylation site to 9-mer motifs (-/+4 residues) | Motifs | UniProt [4] |
| Step 3. Retrieve canonical names of isoform kinases and substrate proteins | Canonical form of isoforms | UniProt [4] |
| Step 4. Retrieve biological links of kinases and substrate proteins | Molecular function (Gene Ontology) | UniProt [4] |
|  | Biological Processes (Gene Ontology) |  |
|  | Cellular Component (Gene Ontology) |  |
|  | Protein Pathways (Reactome) |  |
|  | Protein Tissue of Expression | Human Protein Atlas [5] |
| Step 5. Retrieve links between protein, protein complexes and protein pathways | Protein Complexes | Reactome [6] |
|  | Involvement of protein complexes in protein pathways | Reactome [6] |
| Step 6. Retrieve kinase links | Kinase domains | InterPro [7] |
|  | Kinase homologous superfamily |  |
|  | Kinase families | Zhou et al. [8] |
| Step 7. Retrieve hierarchical relationships | Hierarchical relationships of gene ontology terms | Gene Ontology [9, 10] |
|  | Hierarchical relationships of protein pathways | Reactome [6] |
|  | Hierarchical relationships of kinase domains | InterPro [7] |
| Step 8. Retrieve kinase-favored motifs | Motifs of Serine Threonine kinases | Johnson et al. [11] |
|  | Motifs of Tyrosine kinases | Barir et al.[12] |

**Step 1. Collect, kinases, substrate proteins and phosphorylation sites**

**a) Human phosphosites and substrate(protein)s were extracted from iPTMnet [1], PSP [2] and CPTAC-PDC [3].**

**CPTAC-PDC**

(1) phosphosite*.tmt.csv files from the projects listed in the CPTAC_PDC_mainfest files were retrieved. (Date retrieved - 01/08/2024).
(2) Data listed under the columns, ‘Phosphosite’ was retrieved.
(3) NCBI protein identifiers is mapped to UniProt Accession Numbers and phosphosite at the listed position was validated.
(4) Phosphosites with mismatched residues at the listed position were discarded.

**iPTMnet**

(1) Files, ptm.txt were downloaded from iPTMnet, <https://research.bioinformatics.udel.edu/iptmnet/download> (Date downloaded – 01/17/2024).
(2) From ptm.txt, records for phosphorylation and species of ‘Homo sapiens’ were retrieved.

**PhosphositePlus (PSP)**

(1) From PSP download page, <https://www.phosphosite.org/staticDownloads>, the following two files were downloaded, Kinase_Substrate_Dataset.gz, Phosphorylation_site_dataset.gz. (Date downloaded – 01/22/2024).
(2) Human phosphosites were extracted from the two files.
(3) From the extracted phosphosites, substrate proteins were obtained.

**b) Kinases and substrate(protein)s were extracted from iPTMnet and PSP**

(1) Human kinases were retrieved from ptm.txt downloaded from iPTMnet.
(2) Human kinases were extracted from Kinase_Substrate_Dataset.gz downloaded from PSP.

**Step 2. Convert phosphorylation site to 9-mer motifs (-/+4 residues)**

(1) For all the extracted phosphosites, 9-mer motifs (-/+4 residues) were constructed using protein sequence on UniProt SwissProt database. uniprot_sprot.fasta, uniprot_sprot_varsplic.fasta were downloaded from UniProt ftp site on 01/10/2024. When a protein is not present in the two fasta files, UniProt’s REST API was used to retrieve the sequence information.

**Step 3. Retrieve canonical names of isoform kinases and substrate proteins**

(1) When the kinases or substrate proteins are isoforms, canonical form of the protein is retrieved using the data from UniProt.

**Step 4. Retrieve biological links of kinases and substrate proteins**

(1) UniProt-SwissProt protein data was downloaded on 01/31/2024 from <https://ftp.uniprot.org/pub/databases/uniprot/current_release/knowledgebase/complete/uniprot_sprot.xml.gz>
(2) The XML was parsed to retrieve Gene Ontologies (molecular function, biological processes, cellular component), Reactome pathway information for each protein. For kinases, InterPro kinase domains were also retrieved.
(3) Protein’s tissue of expression were retrieved from the file, normal_tissue.tsv.zip downloaded from HumanProteinAtlas, [https://www.proteinatlas.org/download/normal_tissue.tsv.zip on 02/08/2024](https://www.proteinatlas.org/download/normal_tissue.tsv.zip%20on%2002/08/2024).
(4) The data in the file was parsed to retrieve protein and their tissue of expression at cell type level.

**Step 5. Retrieve links between protein, protein complexes and protein pathways**

(1) Two files, ComplexParticipantsPubMedIdentifiers_human.txt, Complex_2_Pathway_human.txt were downloaded from <https://reactome.org/download/current> on 02/06/2024.
(2) The two files were parsed to retrieve the relations, protein to complex information and complex to pathway information.

**Step 6. Retrieve kinase links**

(1) InterPro entries file was downloaded on 01/24/2024 from <https://ftp.ebi.ac.uk/pub/databases/interpro/current_release/entry.list>. The file was parsed to retrieve protein identifier and InterPro entry types, ‘Homologous_superfamily’ and ‘Domain’.
(2) Kinase families were obtained from the data provided by Zhou et al. [8].

**Step 7. Retrieve hierarchical relationships**

(1) Pathway hierarchies were retrieved using the file, ReactomePathwaysRelation.txt downloaded from <https://reactome.org/download/current> on 02/06/2024.
(2) goa_human.gaf was downloaded from <https://current.geneontology.org/products/pages/downloads.html> on 02/06/2024. The file was parsed to retrieve the go term and the type of term (molecular function, biological process or cellular component). Quick GO API, <https://www.ebi.ac.uk/QuickGO/services/ontology/go/terms/graph>” was used to retrieve the hierarchy of GO terms.
(3) Kinase domain hierarchy was retrieved from the file downloaded from InterPro in Step 6.

**Step 8. Retrieve kinase-favored motifs**

(1) Serine threonine kinase atlas data was downloaded from the article by Johnson et al. [12]. Kinases with ranks 1 through 15 for each motif were retrieved.
(2) Tyrosine kinase – motif information was downloaded from the work by Barir et al. [14]. Records with a percentile score >= 90 were extracted.

The collected data is parsed to create triples of the form, <head | relation | tail>, and integrated to create a knowledge graph. For example, the participation of protein, P61981 in pathway, R-HSA-111447 is represented using the triple, ‘P61981|participating_pathway|R-HSA-111447’. The final knowledge graph thus created is deposited on Zenodo, https://doi.org/10.5281/zenodo.14713589.

**Knowledge graph training and validation**

The total number of triples in the training dataset is 4,881,408. This includes 32 relationship types and 360,098 unique nodes. The hierarchical relationship types are not included as part of the validation dataset. Relationships that include a kinase, substrate(protein) or motif are included in the validation dataset for KGE model hyperparameter optimization (Table 2). As the motif to substrate(protein) relationship is established for our kinase-phosphosite classification, the ‘has_motif’ relationship type that connects a substrate(protein) and motif is not included in the validation dataset

Table 2. Lists all the relationship types. It indicates the presence/absence of the relationship type in the validation dataset. If present, it includes the node types it connects from/to.

| **Relationship type** | **Included in validation dataset?** | **Link includes kinase/substrate(protein)/motif?** |
| --- | --- | --- |
| belongs_to_family | Y | Kinase |
| bio_process | Y | Kinase or substrate(protein) |
| capable_of | N | - |
| capable_of_part_of | N | - |
| cellular_comp | Y | Kinase or substrate(protein) |
| expressed_in | Y | Kinase or substrate(protein) |
| has_domain | Y | Kinase |
| has_motif | N | - |
| homologous_superfamily | Y | Kinase |
| is_a | N | - |
| is_a(domain) | N | - |
| is_a(family) | N | - |
| is_a(form) | N | - |
| k_specific_motif | Y | Kinase, motif |
| mol_func | Y | Kinase or substrate(protein) |
| negatively_regulates | N | - |
| occurs_in | N | - |
| participating_pathway | Y | Kinase or substrate(protein) |
| part_of | N | - |
| part_of_complex | Y | Kinase or substrate(protein) |
| pathway_event_of | N | - |
| positively_regulates | N | - |
| regulates | N | - |
| residue_1 | Y | Motif |
| residue_2 | Y | Motif |
| residue_3 | Y | Motif |
| residue_4 | Y | Motif |
| residue_5 | Y | Motif |
| residue_6 | Y | Motif |
| residue_7 | Y | Motif |
| residue_8 | Y | Motif |
| residue_9 | Y | Motif |

**Protein interaction data retrieval steps**

Protein interaction data is collected from three data sources - BioPlex [13], BioGRID [14], and STRING [15].

**BioGRID**

(1) BioGRID protein interaction data was retrieved from files, BIOGRID-PROJECT-kinome_project_sc-4.4.229, BIOGRID-ALL-4.4.229.tab3 downloaded from <https://downloads.thebiogrid.org/BioGRID/Release-Archive/BIOGRID-4.4.229/> and kinase-specific interaction data was downloaded from paper reported in BioGRID [16]. The two files were downloaded on 01/24/2024.
(2) Interaction data of human proteins were retained. We filtered data from low throughput physical experimental system type.

**BioPlex 3.0**

(1) Protein interaction data was downloaded from <https://bioplex.hms.harvard.edu/data/BioPlex_293T_Network_10K_Dec_2019.tsv> on 01/24/2024.
(2) Data with pW (probability of Wrong) < 0.1 to retrieve high confidence interactants.

**STRING**

(1) Human protein interaction data was downloaded from [https://stringdb-downloads.org/download/protein.links.detailed.v12.0/9606.protein.links.detailed.v12.0.txt.gz](https://stringdb-downloads.org/download/protein.links.detailed.v12.0/9606.protein.links.detailed.v12.0.txt.gz%20on%2001/24/2024) on 01/24/2024.
(2) Data with a confidence score >= 750 was filtered.
