## Supplementary File 2 for "KSMoFinder - Knowledge graph embedding of proteins and motifs for predicting kinases of human phosphosites"

Table 1: Results of including additional features (kinase domain sequences and protein structures) via embeddings from other pre-trained models.

| **Test No.** | **Test Description** | **Features & Embedding sources** | **Testing dataset 1** | | **Testing dataset 2** | |
| --- | --- | --- | --- | --- | --- | --- |
|  |  |  | **ROC-AUC** | **PR-AUC** | **ROC-AUC** | **PR-AUC** |
| 1 | With KSMoFinder-KGE only | Protein biological associations, 9-mer motif (**KSMoFinder-KGE**) | **0.906±0.004** | **0.906±0.006** | **0.93±0.003** | **0.801±0.009** |
| 2 | Inclusion of ProstT5-based protein structure information | Protein biological associations, 9-mer motif (**KSMoFinder-KGE**)  Protein structure (ProstT5) | **0.899±0.006** | **0.888±0.008** | **0.923±0.004** | **0.782±0.004** |
| 3 | Inclusion of Phosformer-based kinase domain sequence and motif sequence information | Protein biological associations, 9-mer motif (**KSMoFinder-KGE**) Kinase domain sequence, 15-mer motif (Phosformer) | **0.898±0.006** | **0.893±0.006** | **0.925±0.003** | **0.785±0.006** |
| 4 | Inclusion of ProstT5 (structure) and Phosformer (kinase domain sequence and motif sequence) information | Protein biological associations, 9-mer motif (**KSMoFinder-KGE**) Kinase domain sequence, 15-mer motif (Phosformer)  Protein structure (ProstT5) | **0.893±0.006** | **0.886±0.008** | **0.92±0.003** | **0.777±0.009** |
| 5 | Effect of dropping KSMoFinder-KGE | Kinase domain sequence, 15-mer motif (Phosformer)  Protein structure (ProstT5) | 0.724±0.004 | 0.652±0.004 | 0.802±0.004 | 0.422±0.004 |
| 6 | Effect of dropping proteins’ biological information contributed via KSMoFinder-KGE | 9-mer motif (KSMoFinder-KGE)  Protein structure (ProstT5) | 0.831±0.005 | 0.794±0.01 | 0.876±0.004 | 0.628±0.01 |

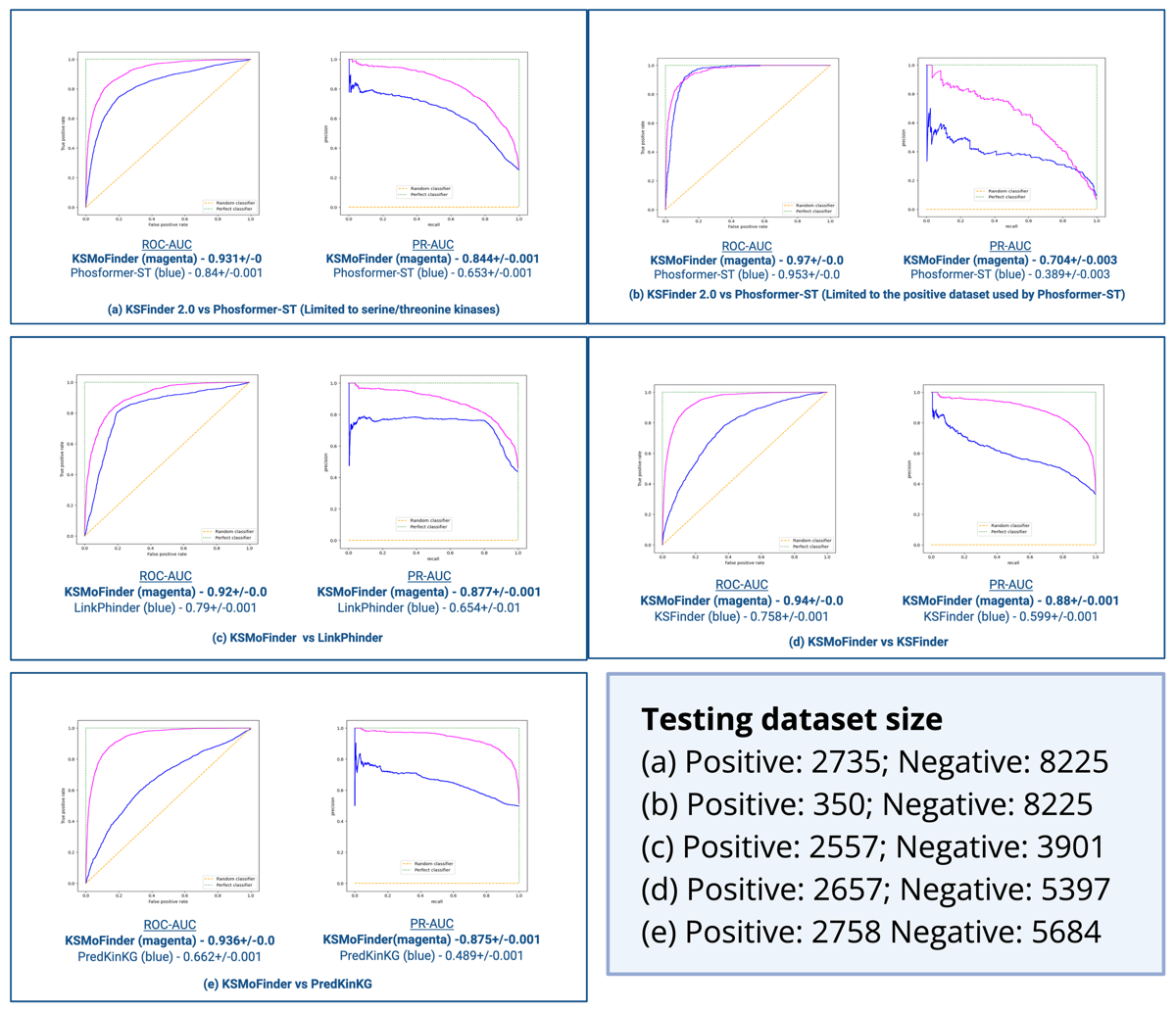

Figure 1. Pairwise comparative evaluation results of KSMoFinder with other models using Testing Dataset 2. Score are reported in ROC-AUC and PR-AUC. (a) KSMoFinder vs Phosformer-ST using the positive dataset from iPTMnet and PSP, (b) KSMoFinder vs Phosformer-ST using the subset containing positive samples reported by Johnson et. al., [4], (c) KSMoFinder vs LinkPhinder, (d) KSMoFinder vs KSFinder, (e) KSMoFinder vs PredKinKG.

Table 2. Comparative evaluation results of KSMoFinder KGE with other protein-based models on testing datasets with and without easy test scenarios.

| **Model** | **Without easy test scenarios** | | | | **With easy test scenarios** | | | |
| --- | --- | --- | --- | --- | --- | --- | --- | --- |
|  | **Testing dataset 1**  (Positive: 2444;  Negative: 2444) | | **Testing dataset 2**  (Positive: 2444;  Negative: 7240) | | **Testing dataset 1**  (Positive: 3025;  Negative: 3025) | | **Testing dataset 2** (Positive: 3025;  Negative: 8489) | |
|  | **ROC-AUC** | **PR-AUC** | **ROC-AUC** | **PR-AUC** | **ROC-AUC** | **PR-AUC** | **ROC-AUC** | **PR-AUC** |
| KSMoFinder | **0.828±0.006** | **0.81±0.007** | **0.899±0.005** | **0.746±0.013** | **0.851±0.008** | **0.839±0.008** | **0.914±0.003** | **0.79±0.009** |
| ProtT5 | 0.714**±**0.01 | 0.686**±**0.012 | 0.851**±**0.05 | 0.61**±**0.014 | 0.752**±**0.007 | 0.726**±**0.001 | 0.871**±**0.04 | 0.676**±**0.009 |
| ESM2 | 0.659**±**0.012 | 0.626**±**0.011 | 0.836**±**0.004 | 0.563**±**0.011 | 0.691**±**0.009 | 0.659**±**0.01 | 0.854**±**0.003 | 0.615**±**0.008 |
| ESM3 | 0.501**±**0.004 | 0.5**±**0.002 | 0.494**±**0.003 | 0.25**±**0.001 | 0.501**±**0.003 | 0.5**±**0.002 | 0.493**±**0.002 | 0.26**±**0.001 |

Table 3. Comparative evaluation results of KSMoFinder and Phosformer-ST (assessment 1) on testing datasets with and without easy test scenarios.

| **KSMoFinder vs Phosformer-ST (sub-assessment 1)** | | | | | | | | |
| --- | --- | --- | --- | --- | --- | --- | --- | --- |
| **Model** | **Without easy test scenarios** | | | | **With easy test scenarios** | | | |
|  | **Testing dataset 1** | | **Testing dataset 2** | | **Testing dataset 1** | | **Testing dataset 2** | |
|  | (Positive: 2276;  Negative: 2276) | | (Positive: 2276;  Negative: 7238) | | (Positive: 2735;  Negative: 2735) | | (Positive: 2735;  Negative: 8225) | |
|  | **ROC-AUC** | **PR-AUC** | **ROC-AUC** | **PR-AUC** | **ROC-AUC** | **PR-AUC** | **ROC-AUC** | **PR-AUC** |
| KSMoFinder | **0.924±0.0** | **0.924±0.001** | **0.924±0.0** | **0.819±0.001** | **0.935±0.0** | **0.937±0.0** | **0.931±0.0** | **0.84±0.001** |
| Phosformer-ST | 0.825±0.001 | 0.818±0.001 | 0.825±0.001 | 0.617±0.001 | 0.837±0.001 | 0.836±0.001 | 0.844±0.001 | 0.653±0.001 |
| **KSMoFinder vs Phosformer-ST (sub-assessment 2)** | | | | | | | | |
| **Model** | **Without easy test scenarios** | | | | **With easy test scenarios** | | | |
|  | **Testing dataset 1** | | **Testing dataset 2** | | **Testing dataset 1** | | **Testing dataset 2** | |
|  | (Positive: 281;  Negative: 281) | | (Positive: 281;  Negative: 7238) | | (Positive: 350;  Negative: 350) | | (Positive: 350;  Negative: 8225) | |
|  | **ROC-AUC** | **PR-AUC** | **ROC-AUC** | **PR-AUC** | **ROC-AUC** | **PR-AUC** | **ROC-AUC** | **PR-AUC** |
| KSMoFinder | **0.962±0.001** | **0.963±0.001** | **0.965±0.001** | **0.656±0.003** | **0.976+0.001** | **0.976+0.001** | **0.97+0.0** | **0.704+0.003** |
| Phosformer-ST | 0.96±0.001 | 0.941±0.002 | 0.949±0.0 | 0.358±0.003 | 0.964±0.001 | 0.948±0.002 | 0.953±0.0 | 0.389±0.003 |
| **KSMoFinder vs LinkPhinder** | | | | | | | | |
| **Model** | **Without easy test scenarios** | | | | **With easy test scenarios** | | | |
|  | **Testing dataset 1** | | **Testing dataset 2** | | **Testing dataset 1** | | **Testing dataset 2** | |
|  | (Positive: 2103;  Negative: 2103) | | (Positive: 2103;  Negative: 3203) | | (Positive: 2557;  Negative: 2557) | | (Positive: 2557;  Negative: 3901) | |
|  | **ROC-AUC** | **PR-AUC** | **ROC-AUC** | **PR-AUC** | **ROC-AUC** | **PR-AUC** | **ROC-AUC** | **PR-AUC** |
| KSMoFinder | **0.898±0.001** | **0.896±0.001** | **0.899±0.007** | **0.859±0.001** | **0.913±0.0** | **0.912±0.001** | **0.912±0.0** | **0.877±0.001** |
| LinkPhinder | 0.798±0.001 | 0.751±0.001 | 0.802±0.001 | 0.672±0.001 | 0.793±0.01 | 0.746±0.001 | 0.79±0.001 | 0.654±0.001 |
