## Supplementary File 3 for "KSMoFinder - Knowledge graph embedding of proteins and motifs for predicting kinases of human phosphosites"

This file contains the hyperparameters used to train the classifiers for different assessments.

The optimal hyperparameters of all the classifiers are determined using a 5-fold stratified cross-validation technique.

Table 1: Lists the final parameters used to train the classifiers for comparative evaluation of embeddings from the four KGE models (Assessment 1).

| **Embedding source** | **Neural Network - hidden layers size** | **Learning Rate** | **Epoch** |
| --- | --- | --- | --- |
| TransE | 40, 50 | 0.01 | 36 |
| DistMult | 40, 50 | 0.01 | 60 |
| ComplEx | 40, 50 | 0.001 | 42 |
| ExpressivE | 40, 50 | 0.001 | 11 |

Table 2: Lists the final parameters used to train the classifiers for comparative evaluation of embeddings from KSMoFinder-KGE and embeddings from external pre-trained models (Assessment 2).

| **Embedding source** | **Neural Network - hidden layers size** | **Learning Rate** | **Epoch** |
| --- | --- | --- | --- |
| KSMoFinder-KGE | 40, 50 | 0.001 | 73 |
| ESM2 | 90, 100 | 0.0001 | 63 |
| ESM3 | 80, 90 | 0.001 | 4 |
| ProtT5 | 80, 90 | 0.001 | 37 |
| Random | 40, 50 | 0.001 | 1 |

Table 3: Lists the final number of epochs used to train the classifiers for assessing the influence of additional feature embeddings from other pre-trained models (Assessment 3). All six classifiers are trained with two hidden layers of size, [40, 50], and a learning rate of 0.001.

| **Test No.** | **Test Description** | **Features & Embedding sources** | **Epoch** |
| --- | --- | --- | --- |
| 1 | With  KSMoFinder-KGE only | Protein biological associations, 9-mer motif (**KSMoFinder-KGE**) | 76 |
| 2 | Inclusion of ProstT5-based protein structure information | Protein biological associations , 9-mer motif (**KSMoFinder-KGE**)  Protein structure (ProstT5) | 36 |
| 3 | Inclusion of Phosformer-based kinase domain sequence and motif sequence information | Protein biological associations, 9-mer motif (**KSMoFinder-KGE**)  Kinase domain sequence, 15-mer motif (Phosformer) | 36 |
| 4 | Inclusion of ProstT5 (structure) and Phosformer (kinase domain sequence and motif sequence) information | Protein biological associations, 9-mer motif (**KSMoFinder-KGE**)  Kinase domain sequence, 15-mer motif (Phosformer)  Protein structure (ProstT5) | 44 |
| 5 | Effect of dropping KSMoFinder-KGE | Kinase domain sequence, 15-mer motif (Phosformer)  Protein structure (ProstT5) | 88 |
| 6 | Effect of dropping proteins’ biological information contributed via KSMoFinder-KGE | 9-mer motif (KSMoFinder-KGE)  Protein structure (ProstT5) | 27 |
